# Stochastic Models for Revealing the Dynamics of the Growth of Small Tumor Populations

**DOI:** 10.1101/743344

**Authors:** Kaitlyn E. Johnson, Amy Brock

## Abstract

The widely accepted model of tumor growth assumes tumors grow exponentially at a constant rate during early tumorigenesis when populations are small. The possibility that tumors might exhibit altered slower growth dynamics or even net cell death below a critical tumor size has yet to be fully explored. Deterministic growth models are capable of describing larger populations because population variation becomes small compared with the average, but when the population being modeled is small, the inherent stochasticity of the birth and death process produces significant variation. Recent advances in high throughput data collection allow for precise and sufficiently large data sets needed to capture this variation. Therefore, we present a stochastic modeling framework to describe and test the potential for altered growth dynamics at small tumor populations.

## I. Introduction to Models of Slowing Cell Growth at Small Populations

The vast majority of studies of cancer cell growth are captured experimentally once populations of cells are large enough that the population average can be feasibly measured, both *in vivo* and *in vitro*. Recent technological advancements allow for the detection of cell growth dynamics at single cell resolution, yet the dynamics that describe this growth regime in cancer are still poorly understood. In the field of ecology, the growth dynamics of a species in an ecosystem has been observed to depend on population density at both high and low density. In cancer, most studies of deviations from exponential growth models have focused on detectable differences at high density only [1]. In this work, we propose incorporating the ecological principle known as the Allee effect to test the relevance of models of slowing growth rates at low population densities [2]. Dynamics of small populations reveals the inherent stochasticity of the birth-death process; thus, we employ stochastic growth models to adequately study the growth dynamics of cancer at early stages of cell growth.

We describe a method of modeling alternate growth hypotheses that incorporate the Allee effect using stochastic growth models. The framework is consistent with that described in [3]. Here, we focus on introducing modifications to the simplest birth-death model that describe the slowing of net growth rate at very low cell numbers by incorporating additional parameters that modulate either the birth or death probability, such as the Master equation in Eq. 1 below.

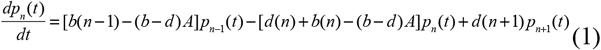

Which describes cells that have a decreased birth probability when N is near the Allee threshold A. From the Master equation, the expected time-evolution of the moments can be derived—here the mean and the variance (Eqs. 2 & 3), as a function of the parameters of interest.

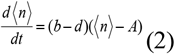

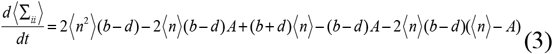

Predicting a behavior in which the average population will go extinct below N (a strong Allee Effect). The relevant outputs of mean and variance in population size N over time can be measured using an *in vitro* experimental workflow in which a large number of low cell density tumor cells are imaged routinely to capture precise growth dynamics of a large number of wells. This data can be combined to obtain the experimentally observed mean and variance of N in time. The stochastic model expected moments of the mean and variance in time can then be used to calibrate to experimentally observed mean and variance to perform parameter estimation using the moment approach [4]. Model selection using Bayesian Information Criterion (BIC) [5] can be used to inform the most parsimonious model structure to describe tumor cells growing at low cell densities and evaluate the relevance of Allee effect models in describing growth at the early stages of tumor cell growth.

## II. Illustrative Results of Stochastic Model Fitting of Allee Growth Dynamics

To illustrate how the stochastic modeling work flow can be applied to experimentally observed growth data, we demonstrate an example study of BT-474 breast cancer cell growth dynamics fit to several stochastic growth models [6]. Data is collected at initial cell numbers of N_0_ = 2, 4, and 10 Fig 1B) and the mean and variance in the cell number N are measured at each time point over a 2-week period (Fig 1C and D). The model is then calibrated to each of seven candidate structural models including a simple birth-death model (the null model), models representing a threshold-like behavior in which N below a threshold A are predicted on average to go extinct (strong Allee effect), and models representing a behavior in which growth is significantly slowed when *n* is near A but still predicted to persist (weak Allee effect). For each of the Allee models, the mechanism of altering growth can either decrease the birth probability, increase the death probability, or affect both, corresponding to differences in expected variances. Based on the BIC values for each model, we find that the mean and variance in time of the BT-474 cells is best described by a weak Allee effect model lowering the birth probability at low N (Fig 1E). Thus, we have evidence that cancer cells growing *in vitro* at low cell densities have a lower birth probability, indicating evidence for the important role of cooperative growth of cancer cells.

**Fig 1.**
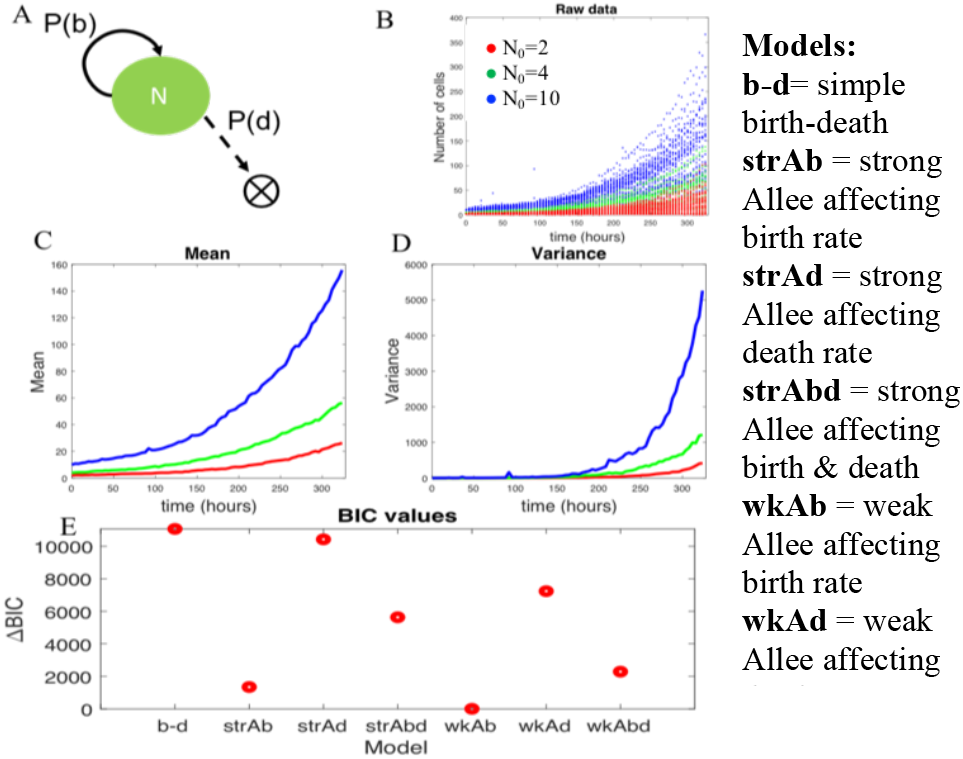
(A) Schematic of stochastic birth-death model, where P(b) denotes the probability of a birth event and P(d) denotes the probability of a death event. (B) Experimentally observed cell number trajectories of BT-474 breast cancer cells at N_0_ = 2, 4, & 10 (red, green blue lines respectively). (C) Mean N in time for each N_0_ (D) Variance in N in time for each N_0_ (E) Model selection results (ΔBIC) after fitting to each stochastic model (abbreviations described on the right) reveals a weak Allee effect model altering the birth rate is the most likely model structure from experimental data.

## III. Quick Guide to the Methods

To implement stochastic modeling methods that incorporate the Allee Effect, the Master equation must be defined using the hypothesis of the nature of the dependencies of stochastic events of birth and death. For example, in the case of a strong Allee effect one could hypothesize that the probability of birth decreases at small N, leading to Eq. 1.

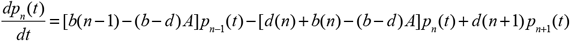

Where *b* represents the inherent birth rate, *d* the inherent death rate, *A* the Allee threshold below which the population goes extinct, and *p_n_* the probability of their being *n* cells at time *t* which gives rise to the resulting distribution. See Table 1 below for a full description of the model parameters in all equations.

**Table 1:**
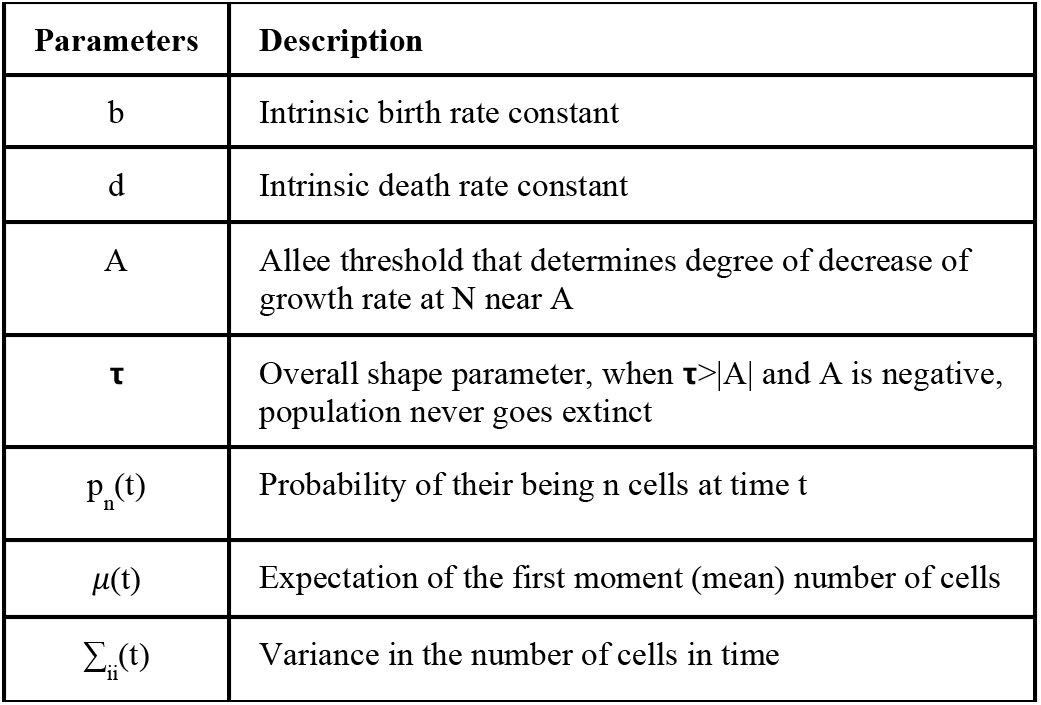
Description of the parameters in each of the equations.

If we apply the Σ*n* and Σ*n*^2^ operator to the master equation, *p_n_*, and using the definition of variance of Σ*_ii_* = 〈*n*^2^〉 〈*n*〉^2^ gives the time derivative of the mean *n* and variance in *n* as a function of the parameters and time as shown in Eqs. 2 & 3.

To generate simulated stochastic trajectories that exhibit these dynamics, one can assume an exponential waiting time distribution and implement the Gillespie algorithm [7] to generate trajectories of cell number in time where birth and death depend on the current population size *n*. We assume that the system is Markovian, i.e. all events occur according to the current population number *n* and a population has no “memory” of its past size evolution. The models presented here assume that the variation in cell number trajectories are due to inherent stochasticity alone, and that all cells are homogenous with the same inherent birth and death probability. In contrast, one could test a stochastic model that results in an observation of slower cell growth at low *n* by implementing more complicated distributions that incorporate two cells types whose birth and death probabilities are dependent on one another, providing a more mechanistic model of interacting cell populations that leads to the observed growth dynamics.

Additional assumptions in parameter estimation from experimental data of cell number in time trajectories assume that the error in the model output, in this case the mean and variance in time, is normally distributed, leading to the likelihood function Eq. 4 below [4]:

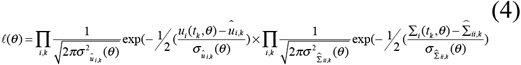

Where u(t, θ) and Σ(t, θ) are the model predicted mean and variance respectively and uk and Σ_k_ are the mean and variance at the kth time point from the data.

### A. Equations

The equations used in stochastic population models are different versions of the Master equation in Eq. 1 and their corresponding moment approximations in Eq. 2 and 3. These correspond to different hypotheses about the mechanism of slowing of growth at low populations. For example, if cell growth follows a weak Allee effect and slows at low populations due to a decrease in birth probability, then Eq. 1, 2, and 3 become:

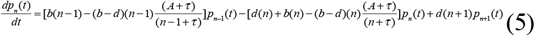

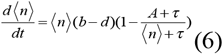

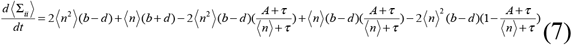

Where Eq. 2 and 3 are derivations of the expected mean and variance from this particular master equation, derived using the methods described in [8] and demonstrated above. Stochastic simulations of a large number of simulations should produce the same mean and variance in time as the model derived approximations of the time evolution of the moments.

### B. Type of settings in which these methods are useful

We are using this method in the setting described to test whether models incorporating a slowing of cell growth at low cell densities are relevant to breast cancer cells growing in *in vitro* experiments. We can apply this method to ask whether other types of cancer cells exhibit early stage growth dynamics that can be described by Allee effect models. The method can also be applied to the in vivo setting to determine whether alternative growth dynamics govern early stage tumor cell dynamics. Stochastic models of this type can be useful for understanding the mechanisms that allow tumor cells to enter the more well-characterized exponential growth phase. An example of a plausible mechanism for an observed Allee effect is that cells require cell-to-cell interactions to cooperatively grow, and these interactions inhibit birth or promote death at low tumor cell densities. The ability to reveal the presence of these altered growth dynamics has implications for understanding how tumors develop and for designing therapeutic strategies aimed at prevent tumor recurrence despite the potential residual presence of circulating tumor cells at a very low cell density.

